# The polysaccharide portion of LPS is critical for caspase-11 inflammasome activation

**DOI:** 10.64898/2026.10.01.756121

**Authors:** Matthew E. Sherman, Colton J. Sheehan, Luca Musante, Igor E. Brodsky, Sunny Shin

## Abstract

Caspase-11 (Casp11) is a murine zymogen that becomes active upon sensing cytosolic lipopolysaccharide (LPS), enabling it to form the non-canonical inflammasome and execute an inflammatory cell death known as pyroptosis. The lipid A acylation state of LPS is thought to be the primary regulator of Casp11 activation. However, our data demonstrate that the polysaccharide moiety of LPS, and particularly the polysaccharide phosphates, are critical for eliciting maximal Casp11 activation. Accordingly, the polysaccharide was required for *Rhodobacter sphaeroides* LPS to act as an inflammasome antagonist. Furthermore, isolated polysaccharide was sufficient to antagonize Casp11 activation. Mechanistically, our findings support a two-step model of non-canonical inflammasome activation where the polysaccharide recruits Casp11 to LPS and the lipid A acylation state governs oligomerization efficiency. These findings provide new insight into the molecular basis of LPS-driven Casp11 activation, with implications for the design of future therapies that target the non-canonical inflammasome.

## Introduction

Lipopolysaccharide (LPS) is a major component of the outer membrane of gram-negative bacteria. While it normally elicits a robust proinflammatory response by innate immune cells that is critical for host defense, when uncontrolled, this inflammatory response can become a primary driver of gram-negative sepsis [1–3]. LPS is composed of a lipid portion, termed lipid A, connected to a polysaccharide moiety of variable length [4]. The lipid A portion consists of a phosphorylated glucosamine backbone connected to acyl chains, whose length and quantity vary across different bacteria [3–6]. The polysaccharide portion is made up of a set of relatively conserved carbohydrates, called the core oligosaccharide, connected to repeating units of serotype-specific sugars, called the O-antigen [7]. Some bacteria lack the enzymes required to attach the O-antigen portion and possess lipooligosaccharide (LOS) instead [8, 9]. Other bacteria lack the core oligosaccharide entirely and possess just lipid A [10–13]. Given that LPS, LOS, and lipid A are all variations of the same molecule, they are collectively referred to as endotoxin.

Endotoxin can be sensed by innate immune cells either on the plasma membrane or within the cytosol. Sensing of endotoxin on the plasma membrane occurs through the Toll-like receptor 4/myeloid differentiation factor 2 (TLR4/MD-2) receptor complex, which robustly induces the expression of pro-inflammatory cytokines [1, 2, 5, 6]. The polysaccharide portion is dispensable for TLR4/MD-2 signaling, as responses are solely dependent on lipid A [14]. Stereotypical lipid A containing 6 acyl chains (hexa-acylated) potently activates TLR4/MD-2 whereas lipid A containing four acyl chains (tetra-acylated) does not [3]. While the structural and biochemical basis for TLR4/MD-2 recognition of endotoxin is well-defined, the basis for cytosolic endotoxin recognition is less understood [15–19].

In murine cells, endotoxin is sensed in the cytosol by the cysteine protease Caspase-11 (Casp11) [20–24]. Casp11 is a zymogen, normally inactive until endotoxin is present, after which it cleaves itself to become active and elicit an immune response [25–27]. To do so, active Casp11 self-oligomerizes into a supramolecular organizing center, termed the non-canonical inflammasome [27–30], where it cleaves and releases the pore-forming subunit of Gasdermin-D (GSDMD) [31–34]. As a result, the cell undergoes a lytic form of cell death called pyroptosis. Prior to complete membrane rupture, potassium effluxes through the GSDMD pore which, in turn, activates the canonical NOD-, LRR-, and pyrin domain-containing protein 3 (NLRP3) inflammasome [35]. The NLRP3 inflammasome activates Caspase-1 (Casp1), which processes IL-1β into its mature form, which is then released from the cell via GSDMD pores [36]. IL-1β is crucial for anti-bacterial defense, but can also contribute to septic shock [37].

Casp11 is thought to respond to the lipid A portion of endotoxin, where the number of acyl chains regulates the extent of activation. Similarly to TLR4/MD-2, hexa-acylated lipid A activates Casp11, whereas most tetra-acylated lipid A structures do not [16, 23, 24]. Furthermore, penta-acylated LPS from *Rhodobacter sphaeroides* (Rs-LPS) which antagonizes TLR4/MD-2 signaling, also antagonizes Casp11 activation [16, 21], highlighting similarities in how Casp11 and TLR4/MD-2 are activated by LPS. However, individual LPS monomers are ligands that bind TLR4/MD-2 [14, 38]. In contrast, in the cytosol, LPS does not exist as a free monomer. Instead, LPS remains either on the bacterial surface, or is released as micellar particles or outer membrane vesicles that enable non-canonical inflammasome formation [17, 18], particularly in regions of positive curvature [19]. This has led us and others [17, 19, 27] to suggest that the mechanisms of cytosolic and cell surface endotoxin sensing are distinct.

Parallels between how TLR4/MD-2 and Casp11 respond to certain lipid A structures have led to a prevailing model in which lipid A acylation is the primary determinant of Casp11 activation. Although the polysaccharide is dispensable for TLR4/MD-2 activation, the role of the polysaccharide in Casp11 activation is unclear. Here, we systematically dissected the contributions of both lipid A and the polysaccharide portion of LPS to non-canonical inflammasome activation by expression or deletion of specific LPS modifying enzymes. Unexpectedly, we found that the presence of a polysaccharide significantly enhanced non-canonical pyroptotic responses, particularly to suboptimal lipid A agonists. Furthermore, this polysaccharide-dependent enhancement was primarily due to phosphates that decorate LPS. We also found that Rs-LPS required the polysaccharide portion to act as a Casp11 antagonist. Moreover, the polysaccharide portion alone was sufficient to antagonize Casp11 responses. Together, our data support a two-step model of non-canonical inflammasome activation, whereby electrostatic interactions between Casp11 and the polysaccharide portion of endotoxin mediate initial recruitment of Casp11, followed by lipid A-directed Casp11 oligomerization. These findings establish the polysaccharide portion of LPS as a crucial mediator of the non-canonical inflammasome response with implications for future anti-sepsis therapeutics.

## Results

### The polysaccharide moiety enhances pyroptosis in response to cytosolic endotoxin

Previous studies from our labs and others have shown that lipid A structure is a key mediator of Casp11-dependent pyroptosis [15, 16, 21, 23, 24]. Accordingly, we modified the lipid A of *Shigella flexneri* 2a LPS (Sf-LPS) to generate structures that have altered acylation or phosphorylation (Fig 1A). To test their effect on Casp11-dependent pyroptosis, we transfected this panel of LPS structures into macrophages and compared their responses to the known unstimulatory Rs-LPS, which contains an atypical lipid A structure (Fig 1B). As expected, unmodified Sf-LPS (Hexa #1) induced robust levels of pyroptosis, whereas Rs-LPS did not (Fig 1C, D). Intriguingly, pyroptosis was largely unaffected by lipid A modification, except for the Tetra #3 variant, whose levels were below that of Hexa #1 but still well above that of Rs-LPS. MALDI-TOF MS validated that the lipid A structures were modified as intended (Fig S1), and ProQ staining of LPS carbohydrates confirmed the polysaccharide was unaffected by lipid A modification (Fig S2A, B). These data indicate that changes in lipid A acylation or phosphorylation have minimal impact on the capacity of Sf-LPS to activate Casp11.

**Figure 1:**
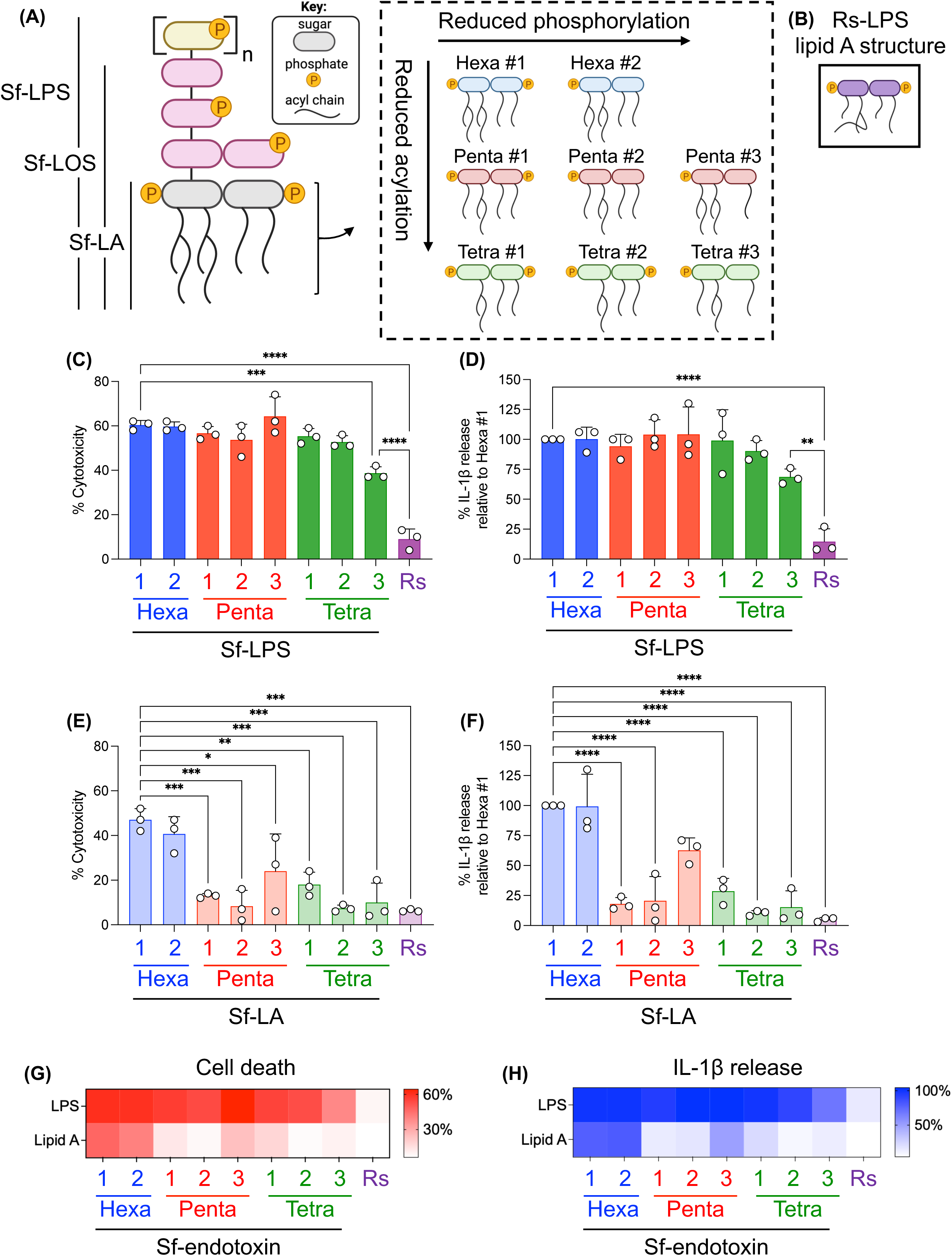
The polysaccharide of LPS enhances pyroptosis. (A) Schematic of *S. flexneri* 2a (Sf) endotoxin with modified lipid A structures. (B) Schematic of the lipid A structure of Rs-LPS. (C) Cell death and (D) IL-1β release 18 hours post-transfection with 1 µg/mL of each type of LPS. Levels of IL-1β were normalized to Hexa #1 Sf-LPS. Raw values are shown in Fig S7. (E-F) Same as in (C-D) except for each type of lipid A. Levels of IL-1β were normalized to Hexa #1 Sf-LA. Raw values are shown in Fig S7. Heat map of (G) cell death and (H) IL-1β release as measured by percent cytotoxicity and percent relative to Hexa #1 Sf-LPS, respectively. Statistical significance determined by ordinary one-way ANOVA where *, **, ***, and **** represent p-values of ≤ 0.05, 0.01, 0.001, and 0.0001, respectively. Dots represent the average values from independent experiments.

In prior studies of how structural changes to LPS acylation impact non-canonical inflammasome assembly, investigators utilized either lipid A, which lacks the polysaccharide entirely, or lipooligosaccharide (LOS), which has a truncated polysaccharide domain [15, 16]. We therefore considered whether the effects of lipid A structural variation on Casp11 activation by Sf-LPS were being buffered by the presence of the polysaccharide. To test this possibility, we isolated the lipid A (Sf-LA) by itself and compared the pyroptotic responses across our entire panel of lipid A structures. Indeed, in the absence of polysaccharide, all the Penta and Tetra Sf-LA structures induced significantly less pyroptosis than Hexa #1 Sf-LA, in many cases down to the levels of Rs-LA (Fig 1E, F). Removal of a single phosphate from lipid A (Hexa #2), however, did not alter the levels of pyroptosis, indicating that lipid A acylation was the primary determinant of pyroptosis in the absence of a polysaccharide, consistent with prior observations [16]. When the entire data set was compared in a heat map format (Fig 1G, H), a marked increase in pyroptosis was observed when the polysaccharide was present on Penta-and Tetra-acylated structures. These data indicate that the presence of the polysaccharide moiety enhances Casp11-mediated pyroptosis.

To determine the region of the polysaccharide responsible for enhanced pyroptosis, we utilized a *galU* mutant of *Sf* which is unable to produce the outer core sugars resulting in LOS instead of LPS [39–42] (Fig S3A), which we validated by ProQ carbohydrate staining (Fig S3B). We engineered this strain to produce tetra-acylated lipid A, which we validated by MALDI-TOF MS (Fig S3C, D). We then compared the pyroptotic responses between LPS, LOS, and lipid A for hexa-and tetra-acylated forms of Sf-endotoxin. Cell death and IL-1β release were significantly decreased in response to hexa-acylated lipid A compared to LPS, and this difference was even more pronounced for tetra-acylated endotoxin (Fig 2A, B). For both hexa-and tetra-acylated endotoxin, LOS induced intermediate levels of cell death and IL-1β release compared to LPS and lipid A. Other markers of pyroptosis, such as GSDMD and Casp1 cleavage, mirrored the differences observed between LPS and lipid A, but in this setting, LOS more closely resembled the phenotype of LPS (Fig 2C).

**Figure 2:**
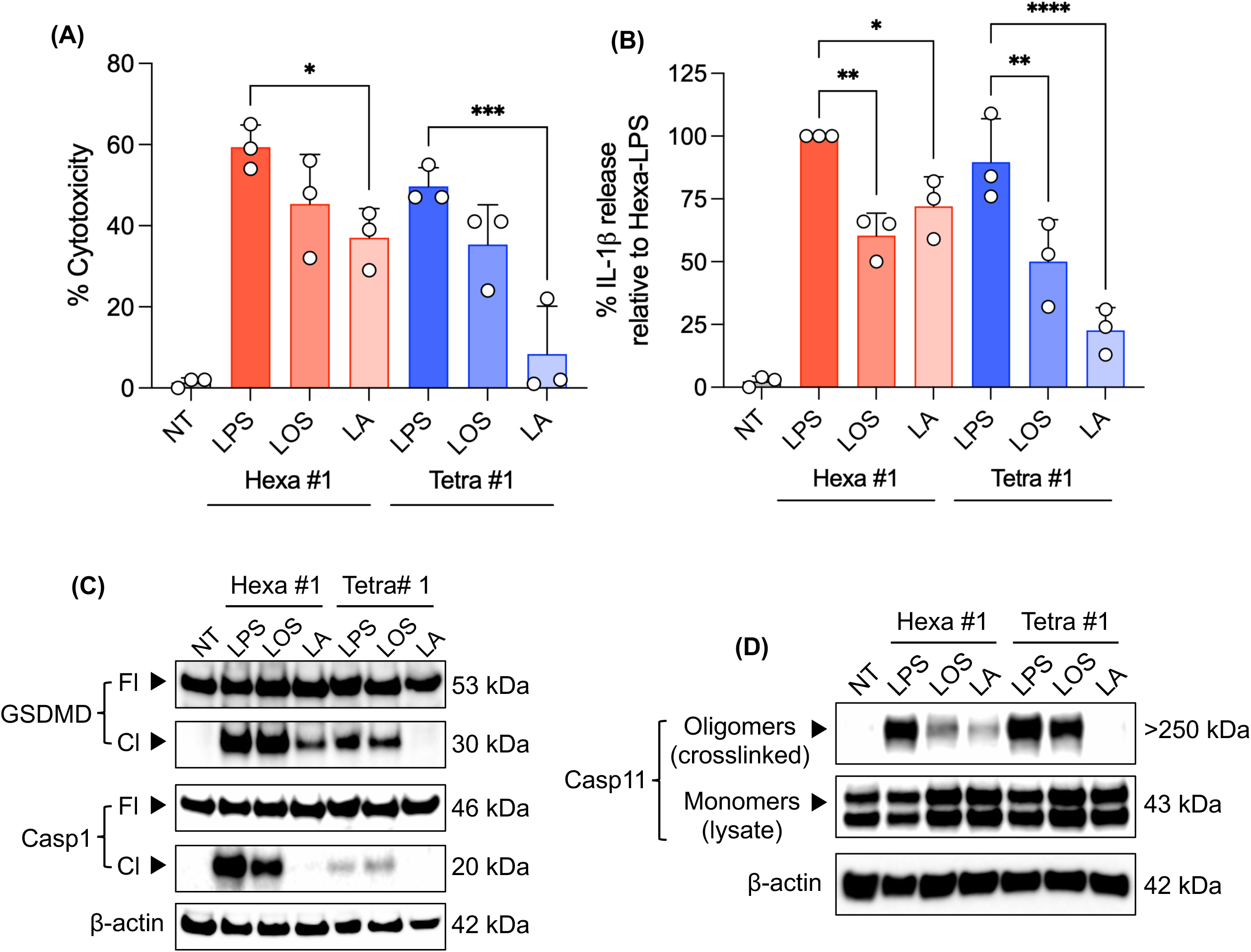
The inner core of the polysaccharide is required for enhanced pyroptosis. (A) Cell death and (B) IL-1β release 18 hours post-transfection with 1 µg/mL of hexa-acylated (Hexa #1) or tetra-acylated (Tetra #1) forms of endotoxin. Levels of IL-1β were normalized to Hexa #1 LPS. Raw values are shown in Fig S8. (C) GSDMD and Casp1 cleavage 6 hours post-transfection under those same conditions. Full-length (Fl) and cleaved (Cl) products were probed for in the cell lysates and concentrated supernatants, respectively. The β-actin loading control was from cell lysates. (D) Casp11 products 18 hours post-transfection under those same conditions. Cell lysates were either crosslinked to probe for oligomers (crosslinked) or not crosslinked as a control to assess Casp11 levels (lysate). The β-actin loading control was from crosslinked lysates. Statistical significance determined by ordinary one-way ANOVA where *, **, ***, and **** represent p-values of ≤ 0.05, 0.01, 0.001, and 0.0001, respectively. Dots represent the average values from independent experiments. NT stands for no transfection.

To investigate LPS-induced Casp11 oligomerization in response to these different forms of endotoxin, we utilized a crosslinking-based approach previously used for other cell death-inducing oligomeric complexes [43–46]. In this assay, we observed that Casp11 oligomers (>250 kDa) form only in response to LPS, which occurs in WT macrophages but not in *Casp11*^-/-^ macrophages (Fig S4). Consistent with our prior observations, hexa-acylated Sf-LA induced less Casp11 oligomerization than its corresponding Sf-LPS, and this difference was more pronounced when tetra-acylated (Fig 2D). Sf-LOS induced Casp11 oligomerization to levels that nearly matched Sf-LPS for tetra-acylated endotoxin but were visibly lower than Sf-LPS for hexa-acylated endotoxin (Fig 2D). Taken together, these data suggest that the inner core portion is sufficient for polysaccharide-driven enhancement of Casp11 activation, but that the full polysaccharide further boosts this response.

Given the evidence that LPS micelles, rather than monomers, may serve as a platform for non-canonical inflammasome oligomerization [17–19], we next asked whether the purified endotoxin used in this reductionist setting was in a monomeric or micellar form. Using dynamic light scattering (DLS) we observed that all the Sf-and Rs-endotoxin were present in the micellar form, and the diameters of those micelles changed upon lipid A modification or the loss of the polysaccharide (Fig S5A-C). These changes in diameter, however, did not correlate with reduced Casp11 responses (Figs 1, 2, and S5), suggesting the size differences have relatively minimal impact of the capacity of endotoxin to activate Casp11. Nonetheless, the presence of endotoxin as micelles in our reductionist system prompted us to wonder whether the negatively-charged polysaccharide was promoting Casp11 oligomerization by initially recruiting it to the endotoxin micelle.

### Phosphates that decorate the polysaccharide are key mediators of pyroptosis

The interaction between Casp11 and LPS depends on positively charged residues within the CARD of Casp11 [21]. Thus, we posited that anionic moieties on the polysaccharide may enhance Casp11 attraction to the endotoxin micelle. Given that the sugars of the inner core [8, 9, 42] and O-antigen [47] of Sf-LPS possess negatively charged phosphates, we sought to determine whether phosphates were responsible for the polysaccharide-dependent enhancement of pyroptosis. We dephosphorylated Sf-and Rs-LPS using hydrofluoric acid (HF) which selectively cleaves phosphoester bonds [48–50], but is not expected to otherwise denature the endotoxin [51–53]. Dephosphorylation was confirmed by MALDI-TOF MS, with Sf-LPS losing all phosphates and Rs-LPS retaining one of the lipid A phosphates (Fig S6A, B). Dephosphorylated (Dephos) Sf-and Rs-LPS remained as micelles, although dephos-Rs-LPS micelles became significantly larger in diameter compared to their phosphorylated counterparts (Fig S6C). The dephos-Sf-LPS induced significantly less cell death and IL-1β release than Sf-LPS with phosphates (Fig 3A, B). Rs-LPS caused low levels of cell death regardless of phosphorylation status but had even further reduced IL-1β release in the absence of phosphates. Importantly, whereas phosphorylated Sf-LPS elicited robust GSDMD cleavage, Casp1 cleavage, and Casp11 oligomerization, dephos-Sf-LPS acted like the unstimulatory Rs-LPS where none of those responses occurred (Fig 3C, D). These data indicate that the phosphates present on Sf-LPS are key drivers of Casp11 activation.

**Figure 3:**
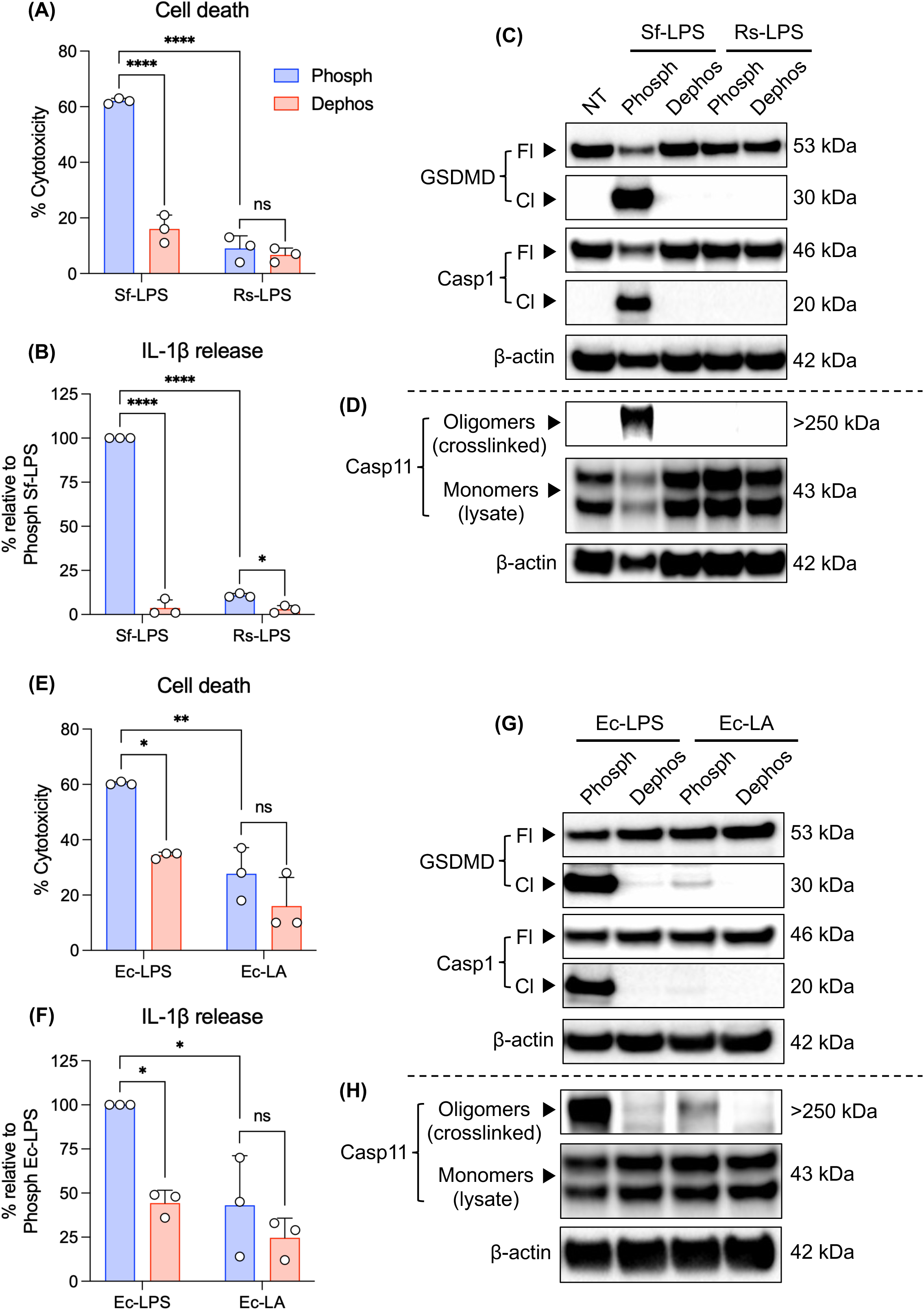
Phosphates the decorate the polysaccharide are key mediators of LPS-induced pyroptosis. (A) Cell death and (B) IL-1β release 18 hours post-transfection with 1 µg/mL of either the phosphorylated (Phosph) or dephosphorylated (Dephos) form of each endotoxin. IL-1β levels were normalized to Phosph Sf-LPS (Hexa #1). Raw values are shown in Fig S9A, B. (C) GSDMD and Casp1 cleavage 6 hours post-transfection under those same conditions. Full-length (Fl) and cleaved (Cl) products were probed for in the cell lysates and concentrated supernatants, respectively. The β-actin loading control was from cell lysates. (D) Casp11 products 18 hours post-transfection under those same conditions. Cell lysates were either crosslinked to probe for oligomers (crosslinked) or not crosslinked as a control to assess Casp11 levels (lysate). The β-actin loading control was from the crosslinked lysates. (E-H) The same as described in (A-D) but for differentially phosphorylated LPS and lipid A from *E. coli* (Ec). IL-1β levels were normalized to Phosph Ec-LPS. Raw values are shown in Fig S9C. Statistical significance determined by 2way ANOVA where *, **, and **** represent p-values of ≤ 0.05, 0.01, and 0.0001, respectively. Dots represent the average values from independent experiments. NT stands for no transfection.

Given that our data suggested the polysaccharide and phosphates are required for maximal pyroptotic responses, we then asked whether these findings extend to the archetypal *E. coli* LPS (Ec-LPS) and its corresponding lipid A (Ec-LA). Consistent with the polysaccharide enhancing pyroptosis induced by Sf-endotoxin, we observed significantly more cell death and increased levels of GSDMD cleavage, Casp1 cleavage, and Casp11 oligomerization in cells transfected with Ec-LPS compared to Ec-LA (Fig 3E-H). To assess the relative contributions of lipid A-associated phosphates compared to polysaccharide-associated phosphates, we dephosphorylated the Ec-LPS and Ec-LA using HF. As seen before, the micellar structure was preserved, and the lipid A retained a single phosphate (Fig S6D-F). Whereas loss of phosphates from Ec-LPS significantly reduced cell death and IL-1β release, only a modest reduction was observed upon loss of phosphates from Ec-LA (Fig 3E, F). The robust GSDMD cleavage, Casp1 cleavage, and Casp11 oligomerization induced by Ec-LPS were abrogated upon dephosphorylation (Fig 3G, H). The relatively lower levels of GSDMD cleavage and Casp11 oligomerization caused by Ec-LA with phosphates were also further reduced upon dephosphorylation. Together, these data indicate that the lipid A phosphates contribute to pyroptosis, though to a lesser extent than the polysaccharide-associated phosphates.

### Rs-LPS requires the polysaccharide to antagonize Casp11 activation

Collectively, our data indicated that the polysaccharide-associated phosphates promote Casp11 activation, suggesting that they contribute to the interaction between LPS and Casp11. Given that Rs-LPS is an antagonist, we next asked whether the polysaccharide is required for its ability to antagonize Casp11 activation by Sf-LPS. We therefore co-transfected Sf-LPS together with 10-fold excess of Rs-LPS or Rs-LA, both of which were unstimulatory in our prior experiments (Fig 1G, H). Rs-LPS was a significantly stronger antagonist of Sf-LPS-mediated cell death, GSDMD cleavage, Casp1 cleavage, and Casp11 inflammasome formation than Rs-LA (Fig 4A-D), indicating that the ability of Rs-LPS to antagonize Casp11 requires its polysaccharide.

**Figure 4:**
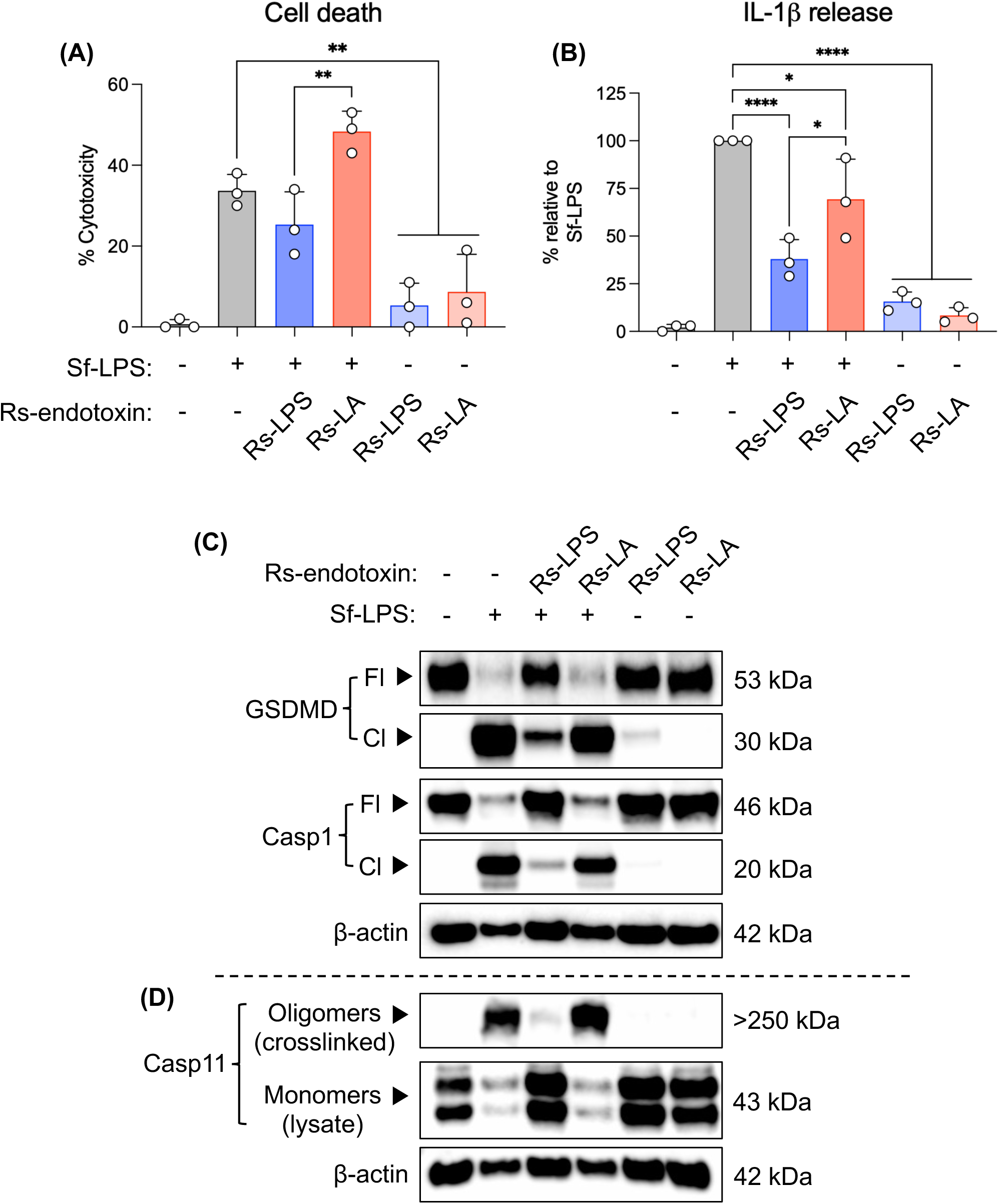
Rs-LPS requires a polysaccharide to be an efficient antagonist. (A) Cell death and (B) IL-1β release 6 hours post-transfection. IL-1β levels were normalized to Sf-LPS (Hexa #1). Raw values are shown in Fig S10. Sf-LPS was transfected at 1 µg/mL whereas *Rs* endotoxin at 10 µg/mL. In the co-transfection setting, Sf-LPS and the indicated *Rs* endotoxin were transfected at the same time. (C) GSDMD and Casp1 cleavage 18 hours post-transfection under those same conditions. Full-length (Fl) and cleaved (Cl) products were probed for in the cell lysates and concentrated supernatants, respectively. The β-actin loading control was from cell lysates. (D) Casp11 products 18 hours post-transfection under those same conditions. Cell lysates were either crosslinked to probe for oligomers (crosslinked) or not crosslinked as a control to assess Casp11 levels (lysate). The β-actin loading control was from the crosslinked lysates. Statistical significance determined by ordinary one-way ANOVA where *, **, and **** represent p-values of ≤ 0.05, 0.01, and 0.0001, respectively. Dots represent the average values from independent experiments.

### Isolated polysaccharide can function as a Casp11 antagonist

Our observation that the polysaccharide is required for optimal efficacy of Rs-LPS to act as a Casp11 antagonist provokes the hypothesis that the polysaccharide portion interacts in some way with the CARD or another portion of Casp11. We therefore asked whether the polysaccharide on its own is sufficient to antagonize Casp11. To address this question, we hydrolyzed the bond in Sf-LPS connecting lipid A to the polysaccharide region and isolated the polysaccharide by size-exclusion chromatography (Fig 5A). ProQ staining of carbohydrates in Sf-LPS compared to the Sf-polysaccharide demonstrated that the native sugar subunit architecture of the polysaccharide was preserved, and importantly, that the dense band below 10 kDa was absent, indicating the removal of lipid A (Fig 5B). Critically, co-transfection of Sf-LPS together with increasing doses of Sf-polysaccharide inhibited cell death and IL-1β release in a dose-dependent manner (Fig 5C, D). Together, these data indicate that the polysaccharide is sufficient to antagonize Casp11 but is insufficient to activate the non-canonical inflammasome in the absence of lipid A.

**Figure 5:**
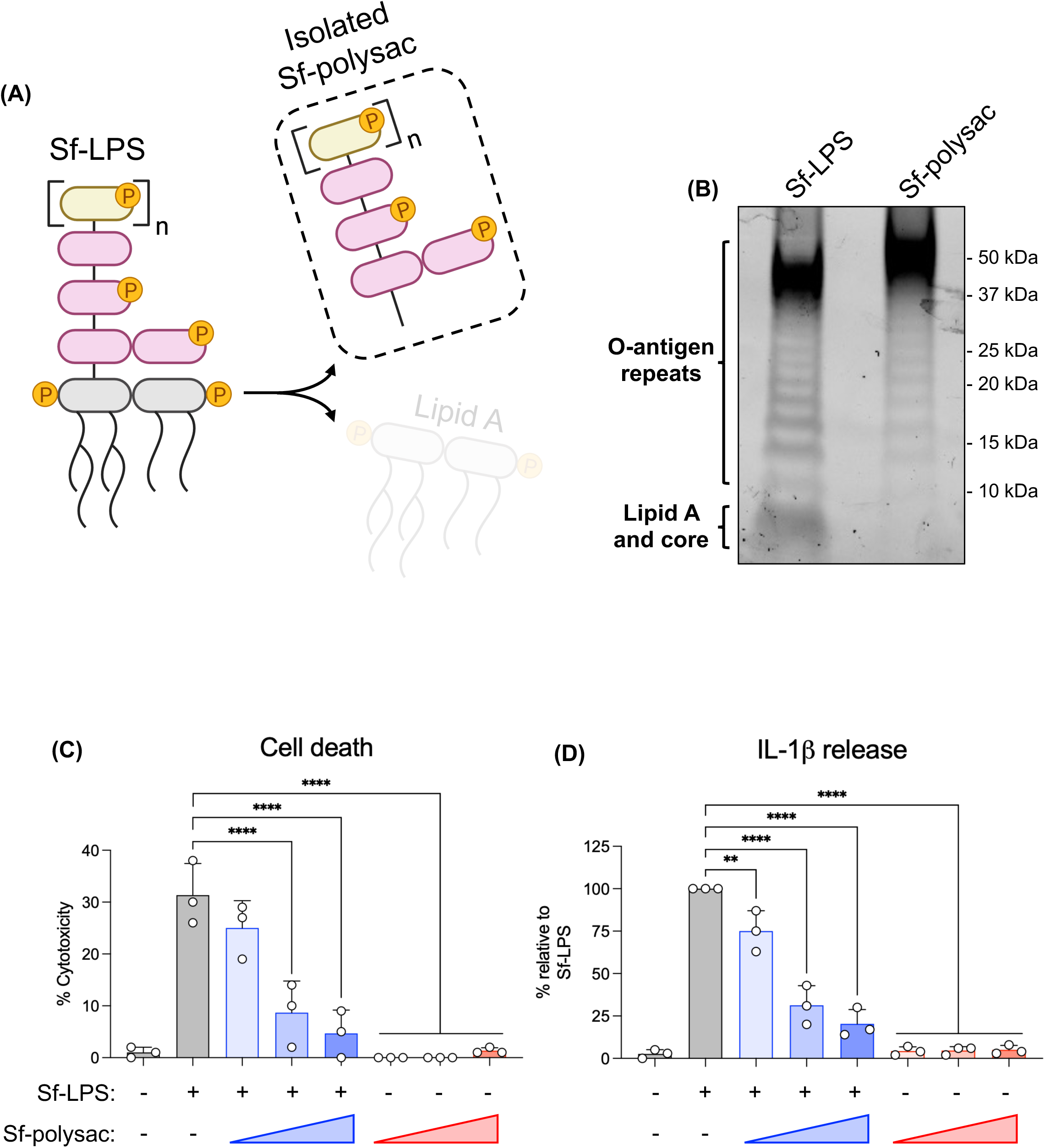
Isolated polysaccharide can function as a Casp11 antagonist. (A) Schematic describing the isolation of Sf-polysaccharide (Sf-polysac) from Sf-LPS. (B) ProQ staining of carbohydrates from Sf-LPS compared to isolated Sf-polysaccharide after separation by SDS-PAGE. Molecular weight designations were made based on the migration of a prestained protein ladder. (C) Cell death and (D) IL-1β release 6 hours post-transfection. IL-1β levels were normalized to Sf-LPS (Hexa #1). Raw values are shown in Fig S11. Sf-LPS was transfected at 1 µg/mL whereas the polysaccharide at increasing doses ranging from 9-36 nM as quantified by Rha sugar content. In the co-transfection setting, Sf-LPS and the Sf-polysaccharide were transfected at the same time. Statistical significance determined by ordinary one-way ANOVA where ** and **** represent p-values of ≤ 0.01 and 0.0001, respectively. Dots represent the average values from independent experiments.

## Discussion

Our data demonstrate that the polysaccharide of LPS mediates optimal non-canonical inflammasome assembly and activation, yet it is insufficient to activate the non-canonical inflammasome without the correct lipid A scaffold. Together, these data support a two-step model in which the polysaccharide and lipid A portions of LPS work in concert to activate the non-canonical inflammasome, most likely at or upon LPS micelles (Fig 6). We propose that Casp11 is initially recruited to endotoxin micelles by electrostatic interactions with the polysaccharide portion, where it then undergoes oligomerization mediated by the acylation status of lipid A. Based on our observations using dephosphorylated endotoxin, we propose that both the polysaccharide and lipid A phosphates mediate the initial recruitment of Casp11 to endotoxin, but that the polysaccharide phosphates are particularly important for optimal recruitment of Casp11. This model provides a basis for understanding why the polysaccharide dramatically enhances Casp11 inflammasome formation and activation, even though lipid A is sufficient to induce Casp11 activation on its own. These findings initially seem contradictory with surface plasmon resonance (SPR) data suggesting Casp11 affinity for lipid A is higher than LPS [21]; however, immobilization of purified Casp11 protein in that setting may not reflect native interactions within the cell. Consistent with our data, numerous studies have reported that pyroptosis is enhanced in response to LPS relative to corresponding lipid A [15, 16, 21, 24], including in the study that performed the SPR analysis [21]. Our study suggests a mechanistic explanation for this interesting but heretofore unclear observation.

**Figure 6:**
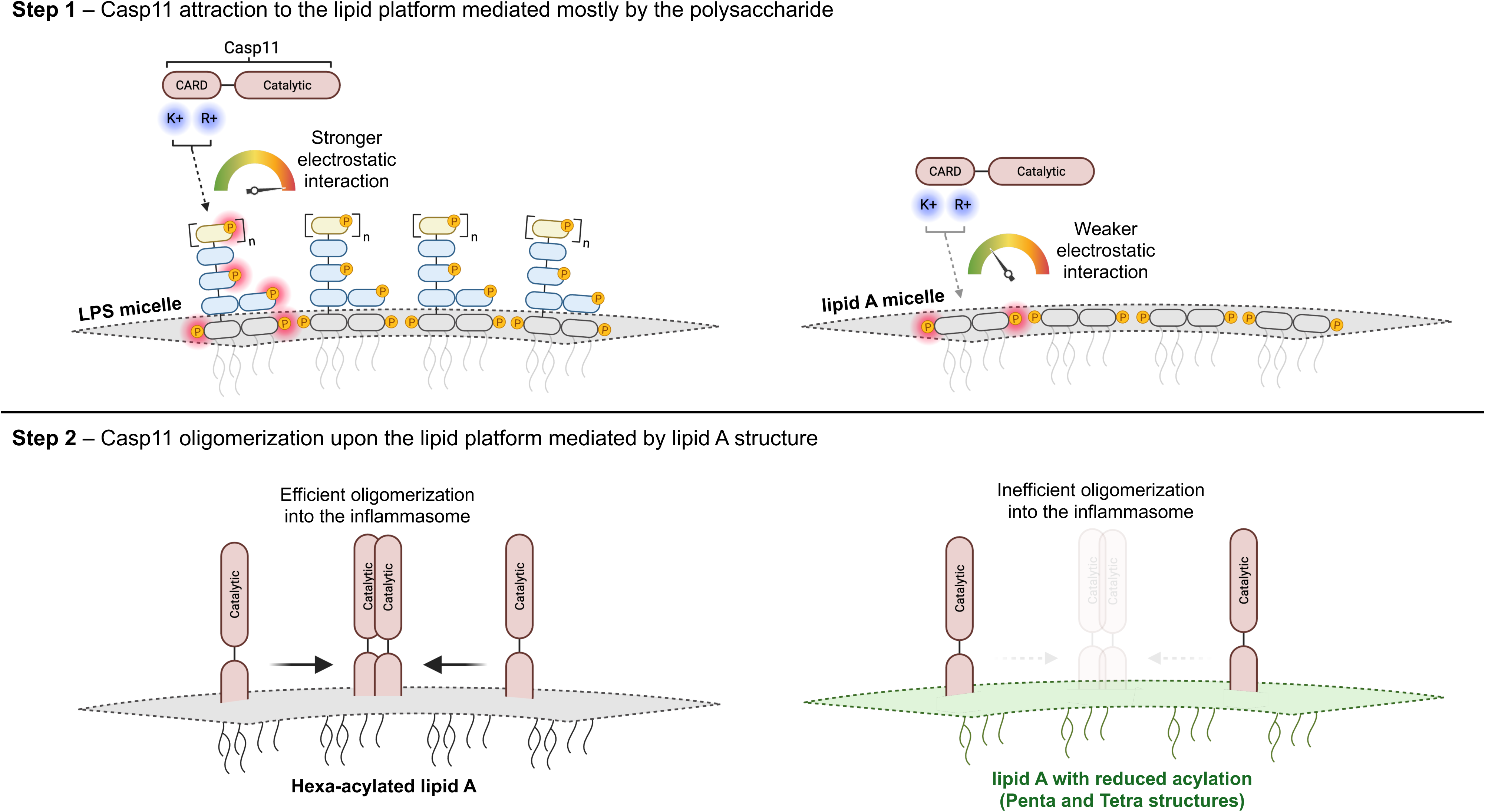
Proposed two-step model of non-canonical inflammasome activation on endotoxin micelles. LPS and lipid A micelles, collectively referred to as endotoxin, serve as lipid platforms for Casp11 oligomerization. First, Casp11 is attracted to the scaffold by electrostatically interacting with phosphates. Attraction to LPS micelles is much stronger than to lipid A micelles since the polysaccharide portion contains more phosphates. Once bound, Casp11 is trafficked along the lipid platform, which depends on the acylation pattern of lipid A. Prototypical hexa-acylated lipid A encourages oligomerization into the inflammasome whereas under-acylated lipid A like Penta and Tetra structures encourage oligomerization less efficiently, or in some cases, not at all.

Our findings that polysaccharide-mediated attraction between Casp11 and endotoxin is insufficient for non-canonical inflammasome formation provide support for a previously perplexing finding that Rs-LPS and prototypical LPS bind Casp11 with equal affinity, yet Rs-LPS does not induce Casp11 activation [21]. Our co-transfection studies comparing Rs-LPS to Rs-LA provide evidence that the polysaccharide of Rs-LPS enables engagement with Casp11, but its atypical lipid A acylation state does not promote Casp11 activation. Polysaccharide engagement with Casp11 is further supported by our observation that Sf-polysaccharide on its own antagonizes Casp11-induced pyroptosis. Polysaccharide-mediated engagement of caspases is also not restricted to Casp11, as Sf-polysaccharide also binds Caspase-3 to inhibit apoptosis [54]. While these findings suggest that the polysaccharide engages with caspases, it is likely that charge-charge interactions with other cationic protein motifs may also occur, raising questions about the potential interactions of polysaccharides more broadly with cellular protein networks.

Inside the cytosol, LPS may not act like ligand, but rather a micellar platform upon which the Casp11 inflammasome forms, as suggested previously by others [17–19]. This may explain why the list of lipid binding partners for Casp11 includes other amphipathic lipids, such as phosphocholines [28, 55, 56], phosphoinositides [28], and cardiolipin [57]. Since these other lipids form micelles, they may also potentially serve as platforms that recruit Casp11. It is possible that, unlike lipid A, these other lipid platforms do not enable Casp11 to undergo its catalysis required for oligomerization [27], however, future biophysical studies are required to interrogate this idea.

Furthermore, the concept that the LPS micellar platform can promote protein oligomerization may extend to other LPS detection systems. The *Limulus polyphemus* (horseshoe crab) complement-like protein factor C is activated by LPS and forms the basis of the *Limulus* amebocyte lysate (LAL) assay. Notably, factor C oligomerizes in response to LPS [58] and disruption of LPS micelles with detergents disables its activity [19, 59]. These findings suggest that factor C may engage LPS micelles via a similar mechanism, whereby its engagement with the LPS micelle enables its oligomerization and activation.

This model of activation may also apply to CASP4 and CASP5, the human orthologs of Casp11. Notably, endotoxin-induced pyroptosis in human macrophages is enhanced by the presence of the core oligosaccharide [15], and CASP4 has been found to binds LPS micelles rather than LPS monomers [17–19]. Given that CASP4 is activated by a broader repertoire of lipid A structures compared to Casp11 [15, 60], it would be of interest to dissect the role of the polysaccharide in promoting CASP4 responses to LPS.

Overall, our findings demonstrate that the polysaccharide of LPS plays a crucial role in Casp11 activation. Uncovering the role of the polysaccharide in noncanonical inflammasome activation improves our understanding of innate immune sensing of LPS, which will aid in designing future therapeutics that abrogate the uncontrolled inflammation that occurs during gram-negative sepsis.

## Materials and Methods

### Reagents and media

Unless otherwise stated, endotoxin free water (Quality Biological #118-325) was used for all assays and stock solutions where applicable. Opti-MEM (Thermo Scientific #31985070) was used for transfections. HBSS (Corning #21-023-CM) supplemented with 10 mM HEPES (Corning #25-060-CI) was used for the SYTOX assay. RPMI-1640 (Corning #10-040-CM) supplemented with 10% fetal bovine serum (FBS, GeminiBio #100-106, Lot#A324069), 10 mM HEPES, 10 mM sodium pyruvate (Corning #25-000-CI), 1% Pen/Strep (P/S, Sigma-Aldrich #P0781), 1 µM β-estradiol (Sigma-Aldrich #E2758), and 10 ng/mL recombinant murine GM-CSF (PeproTech #315-03) was used to maintain Hoxb8 macrophage progenitors. DMEM (Corning #10-017-CV) supplemented with FBS, HEPES, sodium pyruvate, and P/S as described above (β-estradiol and GM-CSF excluded) was used as the base medium for differentiation. For differentiation, L929-cell conditioned supernatant was added to a final concentration of 30%. After differentiation, L929-cell conditioned supernatant added to a final concentration of 10% was used to maintain macrophages in their differentiated state.

### Macrophage sources and procedures

Bone marrow cells from mice were engineered into the macrophage progenitors using the Hoxb8 method as described [61]. Cells were passaged in the progenitor state using the RPMI media containing β-estradiol and GM-CSF described above. To differentiate the progenitors into macrophages, the cells were washed once in PBS to remove residual β-estradiol then seeded in the DMEM media containing 30% L929 supernatant described above for 3 days, with additional media being supplemented 1 day after seeding. Once differentiated, the cells were detached via the cold-PBS method and seeded in the DMEM media containing 10% L929 supernatant described above. In all cases, cells were maintained at 37°C with 5% CO_2_.

### Bacterial strains and genetic manipulation

All strains used in this study were derived from *S. flexneri* 2a 2457T. A double mutant Δ*ospC3*/Δ*galU* was cloned by ƛ-red recombineering [62–64]. Briefly, an *ospC3*::*kan^R^* fragment was PCR amplified from 2457T Δ*ospC3* and subsequently incorporated into the Δ*galU* mutant of 2457T expressing the ƛ-red system. Lipid A expression cassettes, under the control of the *ompC* promoter [65, 66], were cloned into the *attTn7* site by Tn7 transposition [67] as described [68]. These cassettes were incorporated into 2457T Δ*ospC3*, 2457T Δ*msbB1/*Δ*msbB2* double mutant, and 2457T Δ*ospC3*/Δ*galU* double mutant.

### Pro-Q Emerald 300 staining of endotoxin carbohydrates

Endotoxin was extracted from bacterial pellets via the hot aqueous-phenol method as described [69]. Samples were separated via SDS-PAGE using a 14% Tris-Glycine gel (Thermo Scientific #XP00140) fixed overnight in 50% methanol containing 5% acetic acid and stained according to manufacturer’s protocol. When already purified endotoxin or polysaccharide was used, samples were prepared in a buffer of 2% β-mercaptoethanol (BME), 2% sodium dodecyl sulfate (SDS), 10% glycerol, 50 mM Tris-HCl pH 6.8 and <1% bromophenol blue then analyzed as described above.

### Endotoxin purification

LPS was extracted from 2457T strains via the hot phenol-water method [70] and further purified by subsequent extractions to remove contaminating proteins and nucleic acids [71, 72]. Alternatively, LOS was extracted from 2457T Δ*ospC3*/Δ*galU* strains via the phenol-chloroform-petroleum ether method [73]. Lipid A was derived from LPS via the mild acid hydrolysis method [74].

### Commercially purchased endotoxin

Ultrapure LPS from *R. sphaeroides* and ultrapure LPS from *E. coli* O111:B4 were both purchased from Invivogen (#tlrl-prslps and #tlrl-3pelps, respectively).

### Endotoxin procedures

Purified LPS was resuspended at 1 mg/mL in water whereas LOS and lipid A were resuspended at 10 mg/mL in chloroform: methanol (2:1 v/v). When using LOS and lipid A, to avoid contamination of working solutions with chloroform and methanol, the endotoxin was added to an empty tube, dried in a CentriVap centrifugal concentrator (Labconco), then resuspended in the appropriate medium.

### HF dephosphorylation of LPS

All HF work was performed by a trained chemist in a chemical fume hood verified for HF work with all safety precautions carefully adhered to. The methods used herein were adapted from [51, 52]. LPS was resuspended at 10 mg/mL in 49% aqueous HF (Spectrum Chemical Mfg Corp) and incubated for 48 hours at 4°C. Afterwards, the sample was evaporated to dryness under a gentle nitrogen stream for 24 hours. The dried product was resuspended in 10 mL of water, dialyzed against 4 L of water for 24 hours in 1K MWCO tubing, and lyophilized at 0.2 mbar and-50°C in a benchtop freeze dryer (Labconco).

### MALDI-TOF MS analysis of lipid A

Purified lipid A was dissolved in chloroform: methanol (2:1 v/v), deposited on a stainless-steel Bruker MALDI plate, and analyzed as described [68]. To analyze the lipid A structure from purified LPS, we performed a modified version of the FLAT procedure [75, 76]. LPS was first hydrolyzed in a citric acid buffer for 1 hr at 100°C. Then, the lipid A was extracted using chloroform: methanol (2:1 v/v), concentrated by evaporation, and analyzed as described above. Analysis of the lipid A structure from bacterial pellets was also performed as described [77, 78].

### DLS analysis and quantification of endotoxin micelles

Working solutions of endotoxin were diluted in water and analyzed on a ZetaView x30 Nanoparticle Tracking Analyzer (Particle Metrix) with a 488 nm laser wavelength, scatter filter wavelength, laser strength of 80, and shutter of 100. The average particles/mL from three independent injections were used to normalize the exact quantity of endotoxin micelles in each working solution.

### Endotoxin transfection

Macrophages were primed with 1 µg/mL Pam3CSK4 (Invivogen #tlrl-pms) prior to transfection of endotoxin. For 18-hour endotoxin transfections, macrophages were primed for 4 hours. For 6-hour endotoxin transfections, macrophages were primed overnight (∼18 hours). For transfection, the endotoxin was first diluted in opti-MEM so that each sample had equivalent endotoxin particles/mL (400,000 particles/mL). Then FuGENE HD (Promega #E2311) was added to a final concentration of 10% (v/v). The LPS:FuGENE complex was incubated at room temperature for 20 minutes and subsequently added dropwise onto cells at a final concentration of 10,000 particles/mL (representative of approximately 1 µg/mL). Cells were then incubated at 37°C with 5% CO_2_ for the indicated amount time prior to assessing outputs.

### LDH release and SYTOX uptake cytotoxicity assays

At the specified time point after endotoxin transfection, supernatant samples were collected and LDH activity measured by colorimetric assay (Roche #11644793001). The % cytotoxicity was calculated relative to untreated cells and cells completely permeabilized with 1% Triton X-100 (Sigma-Aldrich #X100). To measure plasma membrane pore formation over time, 1 µM SYTOX green nucleic acid stain (Thermo Scientific #S7020) was added to the media and fluorescence measured at 535 nm every 15 minutes using an Infinite 200 Pro plate reader (Tecan #30050303) while cells were maintained at 37°C.

### Standard western blot procedure

Supernatants were collected from treated macrophages, chilled on ice, and precipitated overnight at 4°C by the addition of cold trichloroacetic acid (Sigma-Aldrich #T0699) added to a final concentration of 610 mM. Precipitated proteins were pelleted by centrifugation, washed in cold acetone, and dried at room temperature for 5 minutes. The macrophage cells were lysed in ice-cold RIPA buffer (25 mM Tris-HCl pH 7.6, 150 mM NaCl, 1% NP-40, 1% DOC, 0.1% SDS, and 1X protease inhibitor cocktail) for 5 minutes on ice. Lysate and supernatant samples were mixed with LDS sample buffer (Invitrogen #NP0007) to 1x and DTT to 50 mM. Samples were boiled then separated by SDS-PAGE on a 4-12% Bis-Tris gel (NuPAGE #NP0322BOX) and transferred to a PVDF membrane (Millipore #IPVH00010) via the rapid semi-dry method (1.3 A constant for 7 minutes) using the Trans-Blot Turbo Transfer System (Bio-Rad).

### Modified western blot procedure for inflammasome analysis

Treated macrophages were detached from plates, washed in cold PBS, and lysed in ice cold digitonin buffer (25 mM Bis-Tris HCl pH 7.0, 150 mM NaCl, 5 mM MgCl_2_, 0.02% digitonin) for 20 minutes on ice. Lysates were clarified by centrifugation and crosslinked with freshly prepared disuccinimidyl suberate (DSS, Thermo Scientific #21555) added to a final concentration of 2.5 mM for exactly 30 minutes at room temperature. Afterwards, LDS sample buffer was added to a final concentration of 1x and incubated at room temperature for 15 minutes. Samples were boiled then separated by SDS-PAGE on a 4-12% Bis-Tris gel for 2.5 hours at 150 V surrounded by ice and transferred to a PVDF membrane via the extended semi-dry method (1 A constant for 30 minutes) using the Trans-Blot Turbo Transfer System.

### Antibodies and development of western blots

All membranes were blocked in 5% bovine serum albumin (BSA, Sigma-Aldrich #A7906) and washed with TBST. Membranes were probed with primary antibodies against mouse Casp11 (1:400 dilution, Novus Biologicals #NB120-10454), mouse GSDMD (1:600 dilution, Abcam #AB20985), mouse Casp1 (1:1,000 dilution, Adipogen #AG-20B-0042-c100), or mouse β-actin (1:1,000 dilution, Sigma-Aldrich #A2228). Detection was performed using HRP-conjugated anti-mouse IgG (1:5,000 dilution, Cell Signaling #7076S), anti-rabbit IgG (1:5,000 dilution, Jackson ImmunoResearch #111-035-144), or anti-rat IgG (1:5,000 dilution, Cell Signaling #7077S) and either Pierce ECL substrate solution (Thermo Scientific #32106) or SuperSignal West Femto substrate solution (Thermo Scientific #34095).

### Cytokine ELISAs

Immulon microtiter plates (Thermo Scientific #3455) were used, and each well coated with 1 µg/mL anti-mouse IL-1β antibody (Biolegend #503502) in carbonate-bicarbonate buffer overnight at 4°C. Wells were blocked in PBS containing 1% BSA for at least 1 hour at room temperature and washed in PBST between each subsequent step. Samples, diluted in block buffer, were added to their respective wells and incubated for 2 hours at room temperature. Biotinylated anti-mouse IL-1β antibody (Invitrogen #13-7112-85), diluted to 2 µg/mL in block buffer, was added to each well and incubated for 1 hour at room temperature. SA-HRP (Waters Biosciences #554066), diluted 1,000-fold in block buffer, was added to each well and incubated for 30 minutes at room temperature. O-phenylenediamine (OPD, Sigma-Aldrich #P5412) dissolved in citric acid buffer and supplemented with hydrogen peroxide was used to develop each well into a colorimetric signal which was halted by the addition of sulfuric acid and measured at 490 nm.

### Polysaccharide isolation

A modified approach was adapted from [53, 79]. LPS was hydrolyzed in mild acid hydrolysis buffer as described above. The ethanol wash of the lyophilized product was isolated into a clean tube and evaporated in a CentriVap. The dried product was resuspended in PBS and purified first through a 7K MWCO Zeba size-exclusion desalting spin column (Thermo Scientific #89883) then through a Pierce detergent removal spin column (Thermo Scientific #87777) according to the manufacturer’s guidelines.

### Quantitation of polysaccharide sugar concentration

A modified phenol and sulfuric acid based protocol was adapted from the methods first described [80] and later optimized [81]. Polysaccharide samples or sugar standards were hydrolyzed to monosaccharides in 75% sulfuric acid (v/v) for 30 minutes at room temperature. Then, phenol added to a final concentration of 0.65%, the sample heated at 90°C for 5 minutes, and the colorimetric signal read at 490 nm.

## Acknowledgements

We would like to thank Anh Cao, Pascal Devant, and Jon Kagan for their guidance on Casp11 oligomerization, and particularly Anh Cao, whose advice regarding detergent choice was invaluable. We thank Cammie Lesser and Marcia Goldberg for generously sharing the Δ*msbB1/*Δ*msbB2 and* Δ*galU Shigella* strains and thank Bob Ernst and Jane Michalski for sharing the pSEC10 plasmids used for cloning. We would also like to thank the Mass Spectrometry Core Facility and Extracellular Vesicles Core Facility at the University of Pennsylvania for the use of their resources.

