## Supplemental figures for "The polysaccharide portion of LPS is critical for caspase-11 inflammasome activation"

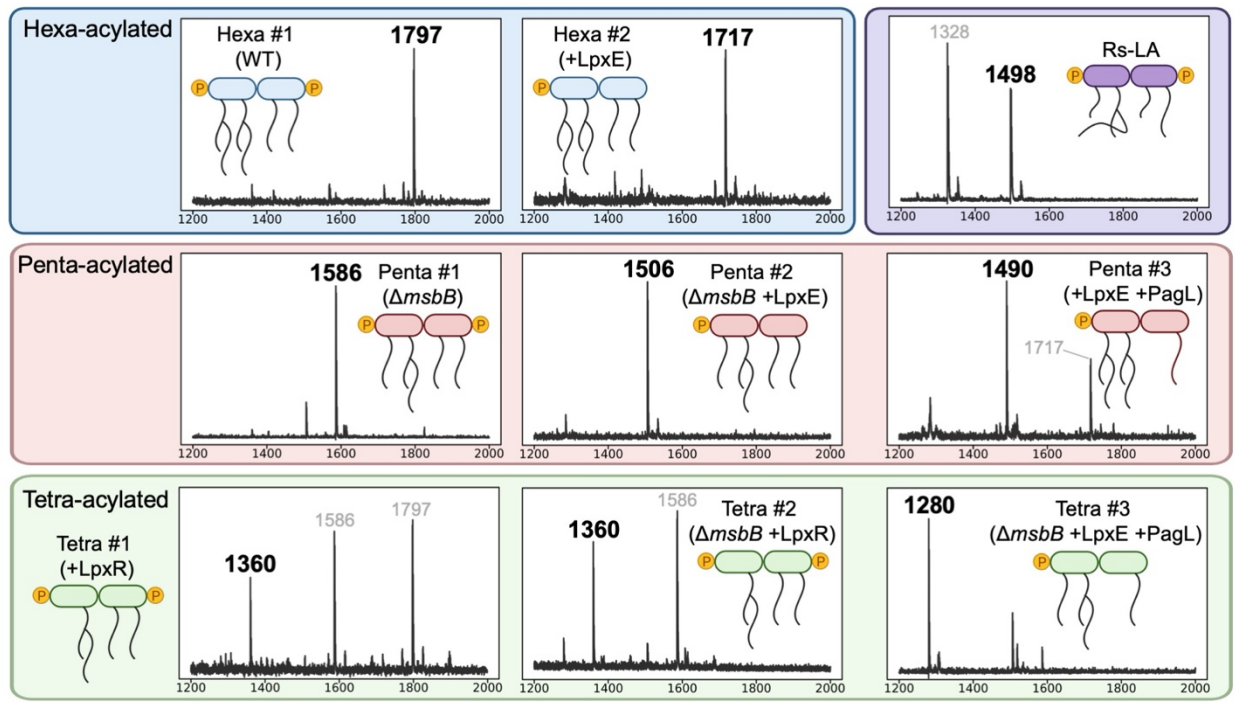

**Figure 1: Lipid A structures were modified as intended.**

MALDI-TOF MS analysis of lipid A samples with relative intensity plotted as a function of the mass to charge ratio ($m/z$). Structures corresponding to the bolded spectral peak are denoted. Other lipid A-related spectral peaks are denoted in grey.

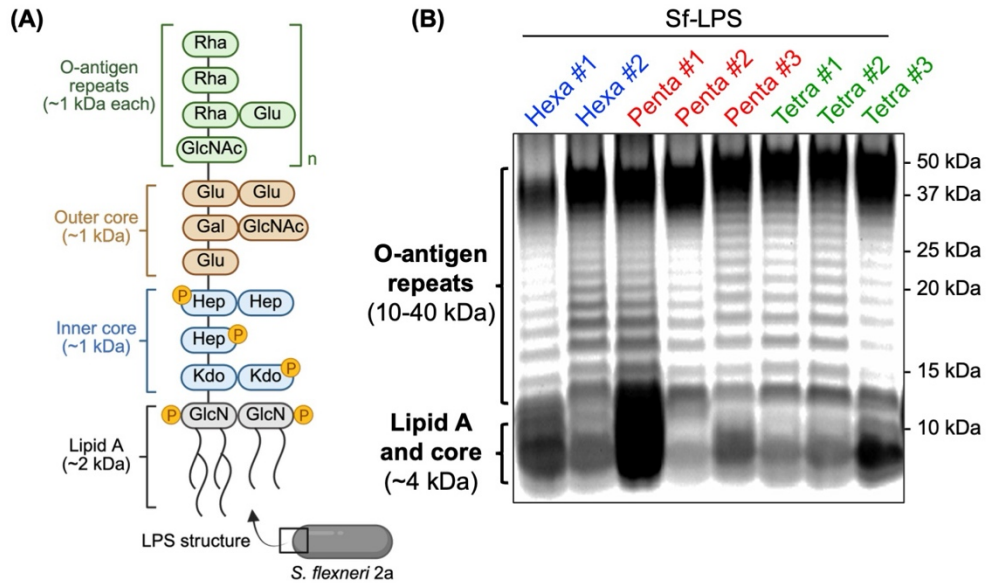

**Figure 2: The polysaccharide was unchanged by lipid A modification.**

(A) The complete structure of *S. flexneri* 2a LPS (Sf-LPS). Sugars are labeled by their abbreviations, acyl chains as the lines drawn on lipid A, and phosphates colored in yellow. Phosphates are predicted to be along the O-antigen but are omitted in this figure due to the uncertainty of their location. (B) ProQ staining of carbohydrates from Sf-LPS extracts after separation by SDS-PAGE.

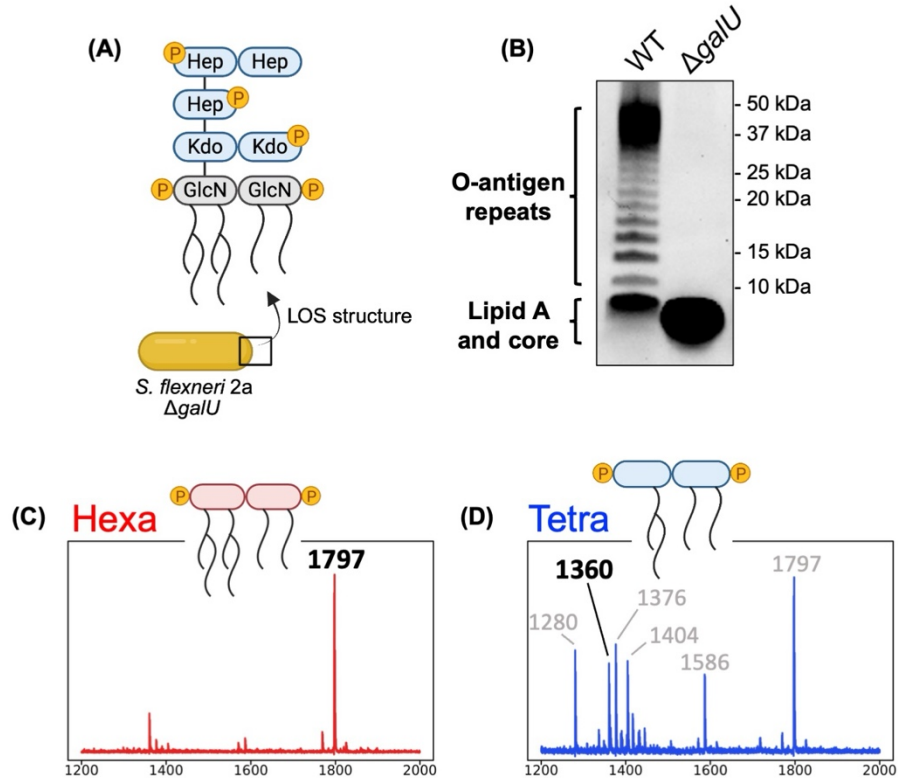

**Figure 3: The *galU* mutant lacks an O-antigen, and the lipid A was modified as intended.**

(A) The complete LOS structure of the *galU* mutant. Sugars are labeled by their abbreviations, acyl chains as the lines drawn on lipid A, and phosphates colored in yellow. (B) ProQ staining of carbohydrates from the LOS extracted from the *galU* mutant compared to LPS extracted from WT *Sf* after separation by SDS-PAGE. (C-D) MALDI-TOF MS analysis of the lipid A portion of the (C) WT *galU* mutant (Hexa #1) and (D) *galU* mutant + LpxR (Tetra #1) with relative intensity plotted as a function of the mass to charge ratio ($m/z$). Structures corresponding to the bolded spectral peak are denoted. Other lipid A-related spectral peaks are denoted in grey.

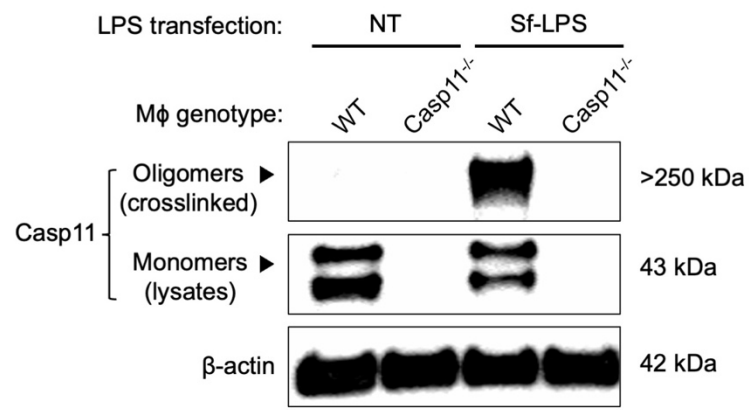

**Figure 4: The crosslinking approach enables assessment of Casp11 oligomers.**

Casp11 products 18 hours post-transfection of WT or Casp11<sup>-/-</sup> macrophages with 1 µg/mL of Sf-LPS (Hexa #1). Cell lysates were either crosslinked to probe for oligomers (crosslinked) or not crosslinked as a control to assess Casp11 levels (lysates). The β-actin loading control was from crosslinked lysates. NT stands for no transfection.

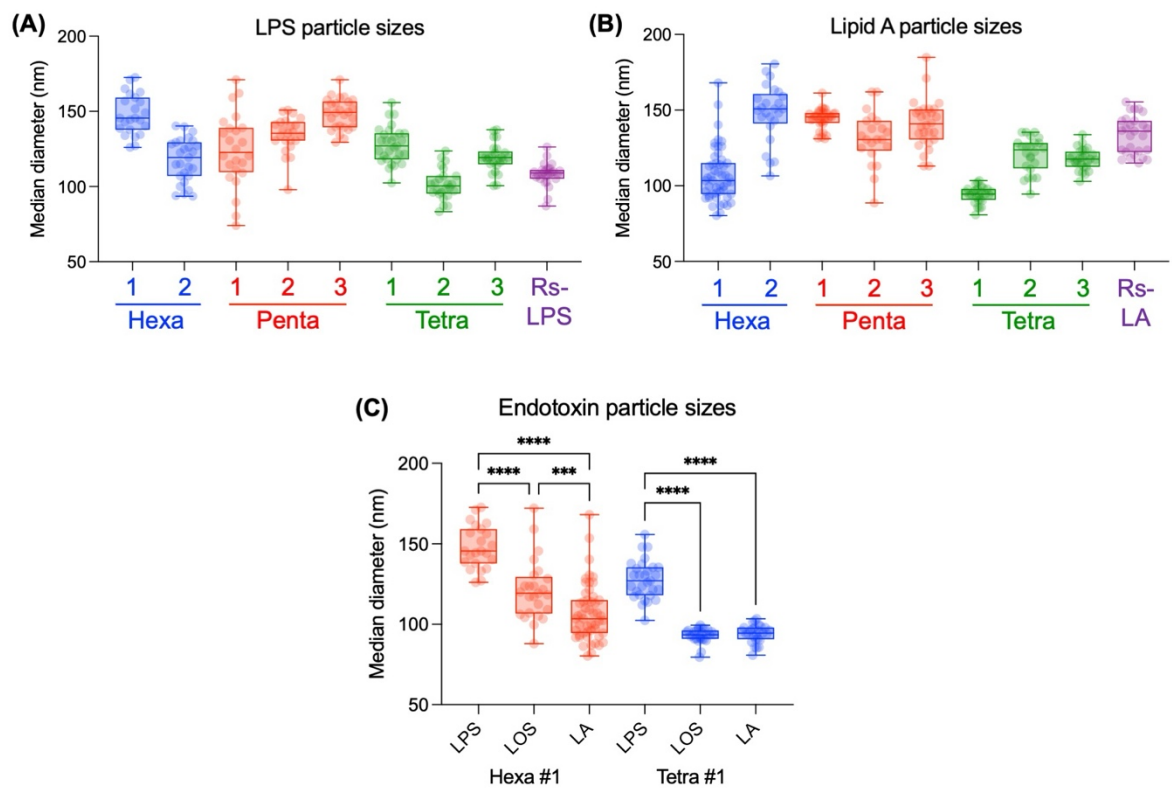

**Figure 5: Endotoxin exists as a micelle whose size varies based on lipid A acylation or the presence of a polysaccharide.**

Endotoxin micelle diameters as measured by DLS for the (A) LPS panel, (B) Lipid A panel, and (C) various forms of Hexa #1 and Tetra #2. Dots represent a read of at least $1 \times 10^6$ particles/mL with at least 20 reads plotted per type of endotoxin. Statistical significance determined by ordinary one-way ANOVA where \*\*\* and \*\*\*\* represent p-values of $\leq 0.001$ and $0.0001$, respectively.

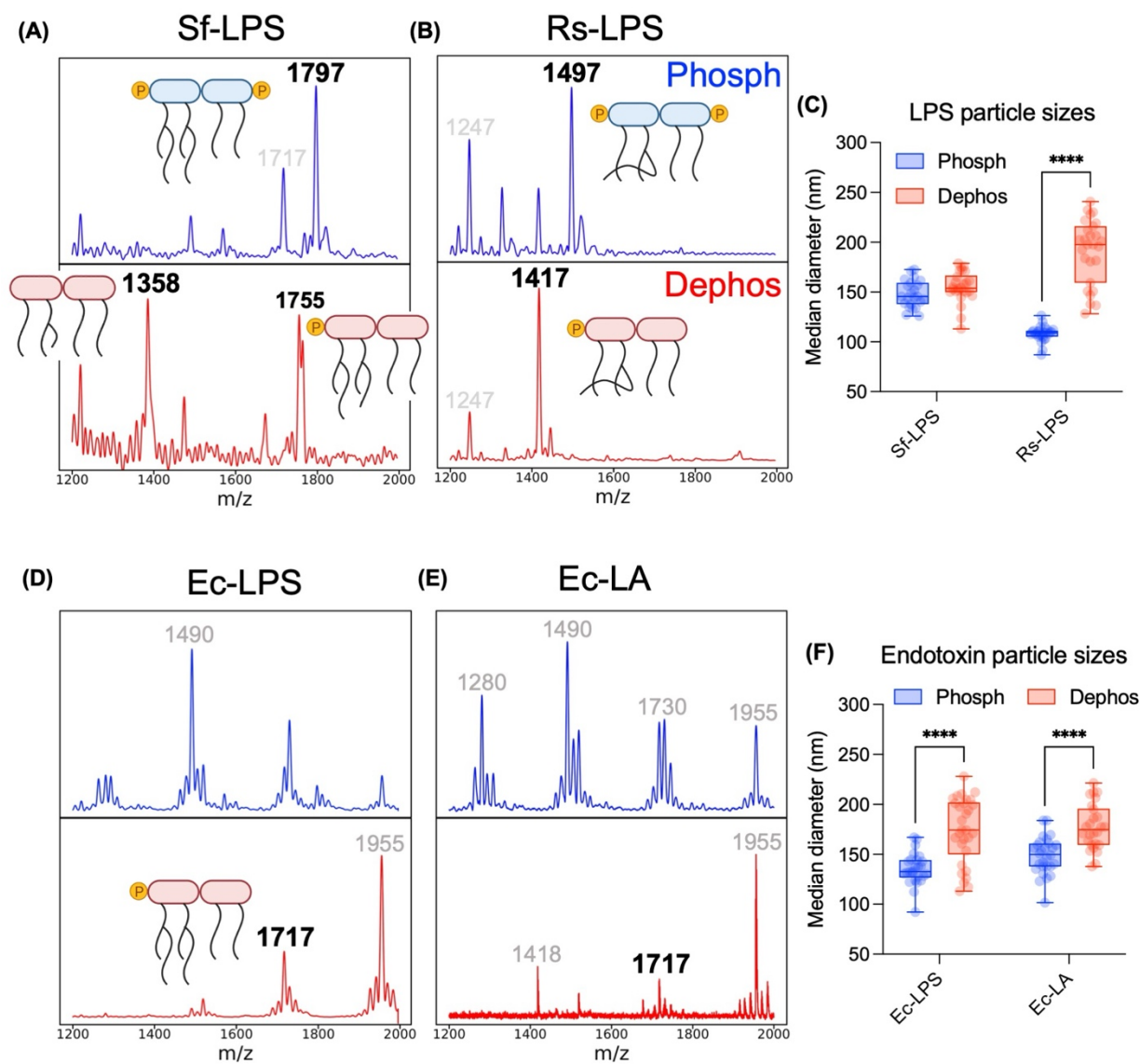

**Figure 6: Validation of the characteristics of dephosphorylated endotoxin.**

MALDI-TOF MS analysis of the lipid A portion of phosphorylated (Phosph) and dephosphorylated (Dephos) (A) Sf-LPS and (B) Rs-LPS with relative intensity plotted as a function of the mass to charge ratio ($m/z$). Structures corresponding to the bolded spectral peak are denoted. Other lipid A-related spectral peaks are denoted in grey. (C) Endotoxin micelle diameters, as measured by DLS, comparing the phosphorylated and dephosphorylated forms of endotoxin. Dots represent a read of at least $1 \times 10^6$ particles/mL with at least 20 reads plotted per type of endotoxin. (D-F) The same as described in (A-C) but for differentially phosphorylated LPS and lipid A from *E. coli* (Ec). Statistical significance determined by 2way ANOVA where \*\*\*\* represents a $p$-value of $\leq 0.0001$.

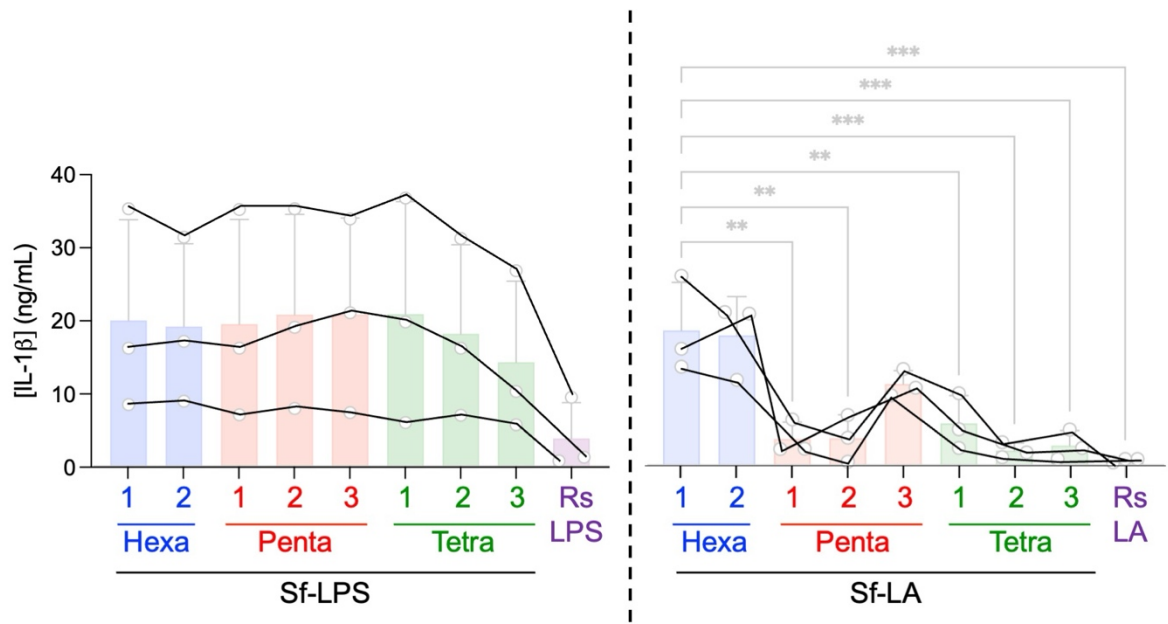

**Figure 7: Plotted values for IL-1β release in response to the LPS and lipid A panel.**

IL-1β release measured 18 hours post-transfection with 1 μg/mL of each type of endotoxin. Statistical significance determined by ordinary one-way ANOVA where \*\* and \*\*\* represent p-values of $\leq 0.05$ and $0.01$ respectively. Dots represent the average values from independent experiments. Lines connect values from the same experiment.

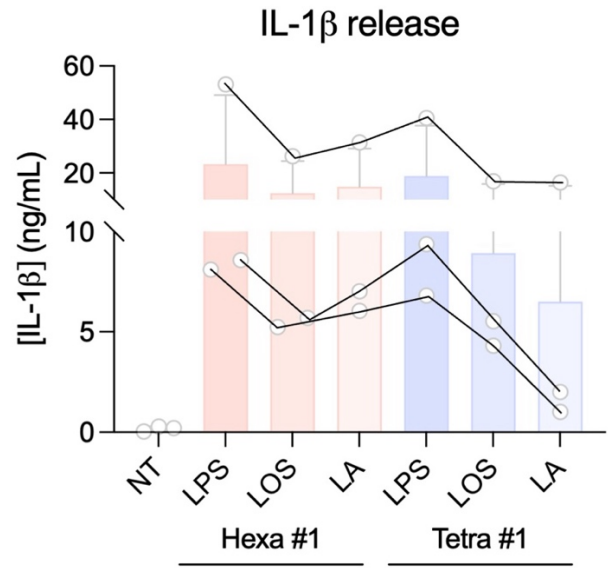

**Figure 8: Plotted values for IL-1β release in response to different forms of endotoxin.**

IL-1β release measured 18 hours post-transfection with 1 μg/mL of either hexa-acylated (Hexa #1) or tetra-acylated (Tetra #1) forms of endotoxin. Dots represent the average values from independent experiments. Lines connect values from the same experiment.

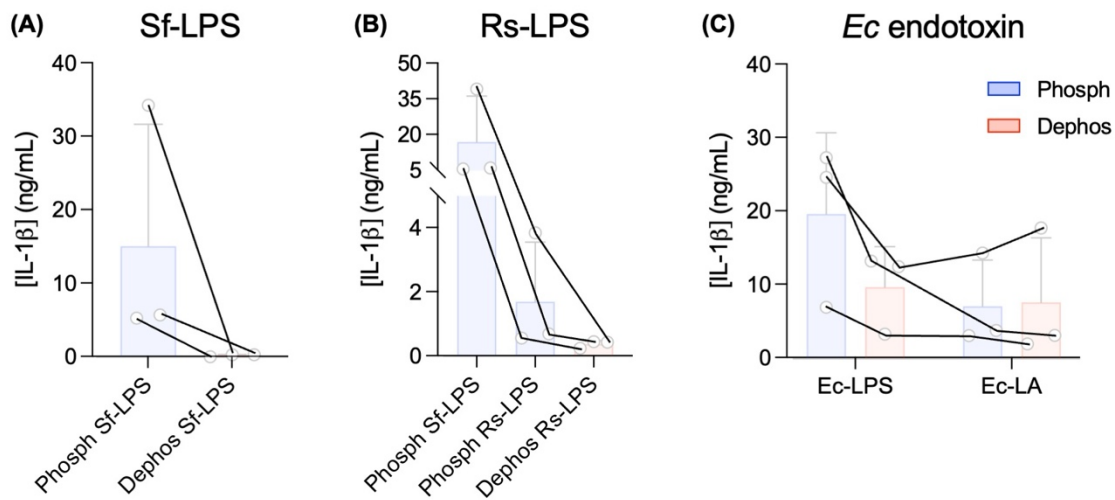

**Figure 9: Plotted values for IL-1β release in response to differentially phosphorylated LPS.**

IL-1β release measured 18 hours post-transfection with 1 μg/mL of either phosphorylated (Phosph) or dephosphorylated (Dephos) forms of endotoxin. (A) Sf-LPS (Hexa #1) compared to Dephos Sf-LPS. (B) Rs-LPS and dephosphorylated Rs-LPS compared to Sf-LPS. (C) Differentially phosphorylated Ec-LPS compared to Ec-LA. Dots represent the average values from independent experiments. Lines connect values from the same experiment.

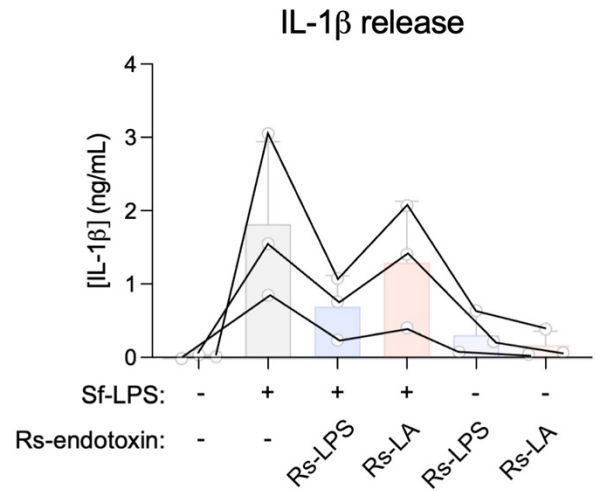

**Figure 10: Plotted values for IL-1β release in response to co-transfection of Sf-LPS with Rs endotoxin**

IL-1β release measured 6 hours post-transfection with 1 μg/mL of Sf-LPS, 10 μg/mL of the indicated form of *Rs* endotoxin, or co-transfection at those indicated doses. Dots represent the average values from independent experiments. Lines connect values from the same experiment.

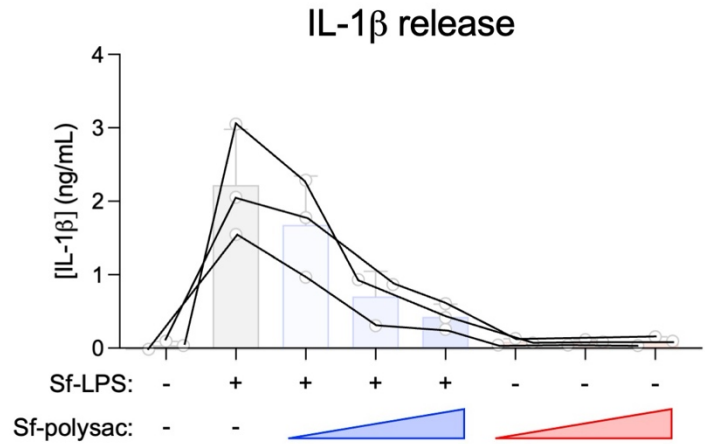

**Figure 11: Plotted values for IL-1β release in response to co-transfection of Sf-LPS with increasing doses of the Sf-polysaccharide.**

IL-1β release measured 6 hours post-transfection with 1 μg/mL of Sf-LPS, 9-36 nM of Sf-polysac, or co-transfection at those indicated doses. Dots represent the average values from independent experiments. Lines connect values from the same experiment.
